# Decreased thermal tolerance as a trade-off of antibiotic resistance

**DOI:** 10.1101/2021.04.05.438396

**Authors:** Cristina M. Herren, Michael Baym

## Abstract

Evolutionary theory predicts that adaptations, including antibiotic resistance, should come with associated fitness costs; yet, many resistance mutations seemingly contradict this prediction by inducing no growth rate deficit. However, most growth assays comparing sensitive and resistant strains have been performed under a narrow range of environmental conditions, which do not reflect the variety of contexts that a pathogenic bacterium might encounter when causing infection. We hypothesized that reduced niche breadth, defined as diminished growth across a diversity of environments, can be a cost of antibiotic resistance. Specifically, we test whether chloramphenicol-resistant *Escherichia coli* incur disproportionate growth deficits in novel thermal conditions. Here we show that chloramphenicol-resistant bacteria have greater fitness costs at novel temperatures than their antibiotic-sensitive ancestors. In several cases, we observed no resistance cost in growth rate at the historic temperature but saw diminished growth at warmer and colder temperatures. These results were consistent across various genetic mechanisms of resistance. Thus, we propose that decreased thermal niche breadth is an under-documented fitness cost of antibiotic resistance. Furthermore, these results demonstrate that the cost of antibiotic resistance shifts rapidly as the environment changes; these context-dependent resistance costs should select for the rapid gain and loss of resistance as an evolutionary strategy.

## Introduction

The extensive distribution of antibiotic resistant bacteria across both clinical and environmental habitats demonstrates the difficulty of limiting the dispersion of antibiotic resistance (Chatterjee et al. 2018). However, evolutionary theory predicts that resistance should come with widespread fitness costs in the absence of antibiotics: if there were no cost, then all pathogens should become resistant, and resistance should never be lost (Law 1979, Luciani et al. 2009, Andersson and Hughes 2010). Empirically, many known resistance mechanisms are energetically costly (e.g. efflux pumps), which divert a portion of a cell’s energetic budget away from growth and reproduction (Wood and Cluzel 2012, El Meouche and Dunlop, 2018). Many studies have sought to quantify fitness costs to antibiotic resistance with the goal of identifying growth deficits that accompany resistance (as reviewed by Durão et al. 2018). Meta-analyses of bacterial growth rates have found wide ranges of fitness costs for resistance to different antibiotics (Melnyk et al. 2015). And, in some cases, there are no discernable growth rate difference between antibiotic-sensitive strains and resistant strains (Sander et al. 2002, Gagneux et al. 2006, Lin et al. 2018). However, an important consideration is that most experimental tests of bacterial growth are done under tightly constrained environmental conditions, usually at 37C to mimic human body temperature. The absence of fitness cost in this single context does not preclude the presence of fitness costs in different conditions (Olivares et al. 2014), meaning there may be unidentified fitness costs to these seemingly unaffected resistant bacteria.

Although bacterial cultures in the laboratory are maintained under a narrow range of environmental parameters, bacterial pathogens might encounter many different environments when causing an infection (Runkel et al. 2013, Fang et al. 2016). For instance, their host’s body temperature might rise or fall (Hasday et al. 2000), or the cell may have to persist outside the body during a transmission event (Berger et al. 2010). Fitness costs to antibiotic resistance might manifest differently in these varied environments; for example, temperature mediates both antibiotic resistance and cell growth by affecting processes such as protein folding and the rate of chemical reactions within the cell (Wilson 2014, Mondal et al. 2014). Thus, a mutation that confers resistance might be neutral at one temperature but could have adverse effects when protein shape or enzyme kinetics are altered. In this case, strong specialization for antibiotic resistance might have costs in the ability to tolerate multiple environments.

Here, we propose decreased thermal niche breadth, defined as reduced growth rate across a range of temperatures, as a cost of antibiotic resistance. Because niche breadth is independent of maximum growth rate, it is possible for fitness costs to manifest as either a cost in maximum growth rate (Fig. 1, left), a cost in thermal niche breadth (Fig. 1, center), or a cost in both maximum growth rate and thermal niche breadth (Fig. 1, right). The addition of this dimension of fitness could reconcile the theoretical framework of requisite fitness costs with the experimental evidence that fitness costs are absent in some conditions; if costs to resistance are in decreased niche breadth, there may be no apparent fitness costs in some environments.

**Figure 1:**
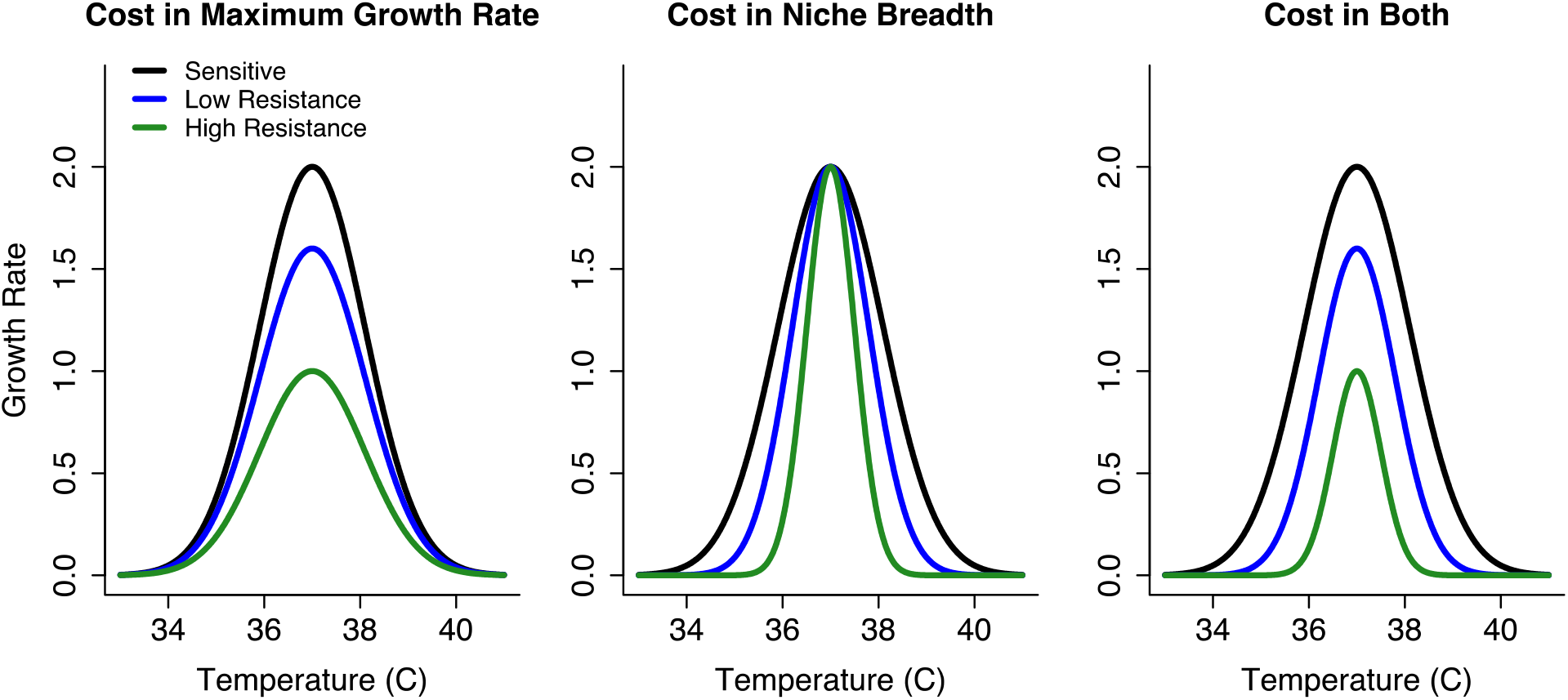
Conceptual illustration of niche breadth and maximum growth rate as independent dimensions of fitness. Because fitness costs in thermal niche breadth are independent of absolute growth rate, increasing resistance might show a decreased growth rate across all temperatures (cost in maximum growth rate; left), a more rapid drop-off in growth rate away from the thermal optimum (cost in thermal niche breadth; center), or both a decreased maximum growth rate and a narrower thermal range (cost in both maximum growth rate and niche breadth; right).

The interest in quantifying fitness costs is, in large part, due to the influence of fitness costs on the population dynamics of antibiotic-resistant and antibiotic-sensitive bacterial strains (Levin et al. 1997, Mortimer et al. 2020). A fitness cost to antibiotic resistance would slow the spread of resistant strain, as the resistant bacteria would be at a competitive disadvantage (Andersson and Hughes 2010). However, it is not straightforward to extrapolate from fitness costs to community composition; several additional dimensions of fitness such as lag time (Adkar et al. 2017), resource use efficiency (Hodapp et al. 2019), and resource storage (de Mazancourt and Schwartz, 2010) also affect competitive outcomes. Thus, while a fitness costs in growth rate represents a deficit in the ability of a strain to quickly reproduce, it is difficult to predict the success of an antibiotic-resistant strain from only these measurements.

In this study, we test the hypothesis that increasing resistance to chloramphenicol carries fitness costs in thermal niche breadth, defined as growth rate across multiple temperatures. We selected chloramphenicol as the focal drug due to its many resistance pathways, which allows us to explore the consistency of fitness costs across a variety of genetic mechanisms of resistance (Crofts et al. 2019). In addition, chloramphenicol resistance spans a large gradient, where evolved MICs can be in excess of 100-fold the initial MIC (Toprak et al. 2012). We evolved resistant bacterial populations by conducting 24 parallel evolution experiments, beginning from a culture of *Escherichia coli* from the Keio collection. We measured the growth rates of these strains at novel temperatures to evaluate how resistance affected thermal tolerance. Additionally, we competed strains of varying resistance against one another to investigate how fitness costs of resistance translate to community composition and the competitive success of highly resistant strains. Finally, we sequenced the genomes of isolates from each lineage to understand the genetic basis of resistance and to test whether mutations in specific genes had differential fitness effects in novel temperatures. Together, this study defines thermal niche breadth as a dimension of fitness, quantifies how these fitness costs shape the competitive success of resistant strains, and relates these deficits in novel environments to the genomic basis of resistance.

## Results

### Obtaining bacterial strains with varied resistance levels

We experimentally evolved 24 replicate lineages of *E. coli* to tolerate increasing concentrations of chloramphenicol. By serially passaging bacterial cultures through 14 increasing chloramphenicol levels, we obtained 336 (24 lineages x 14 concentrations) populations of *E. coli* across a gradient of resistance levels (Fig. 2). The 24 replicate lineages enabled us to study the variability arising from the stochastic nature of mutation acquisition. We refer to these populations as “cultures” rather than “strains” due to the possible coexistence of multiple genotypes.

**Figure 2:**
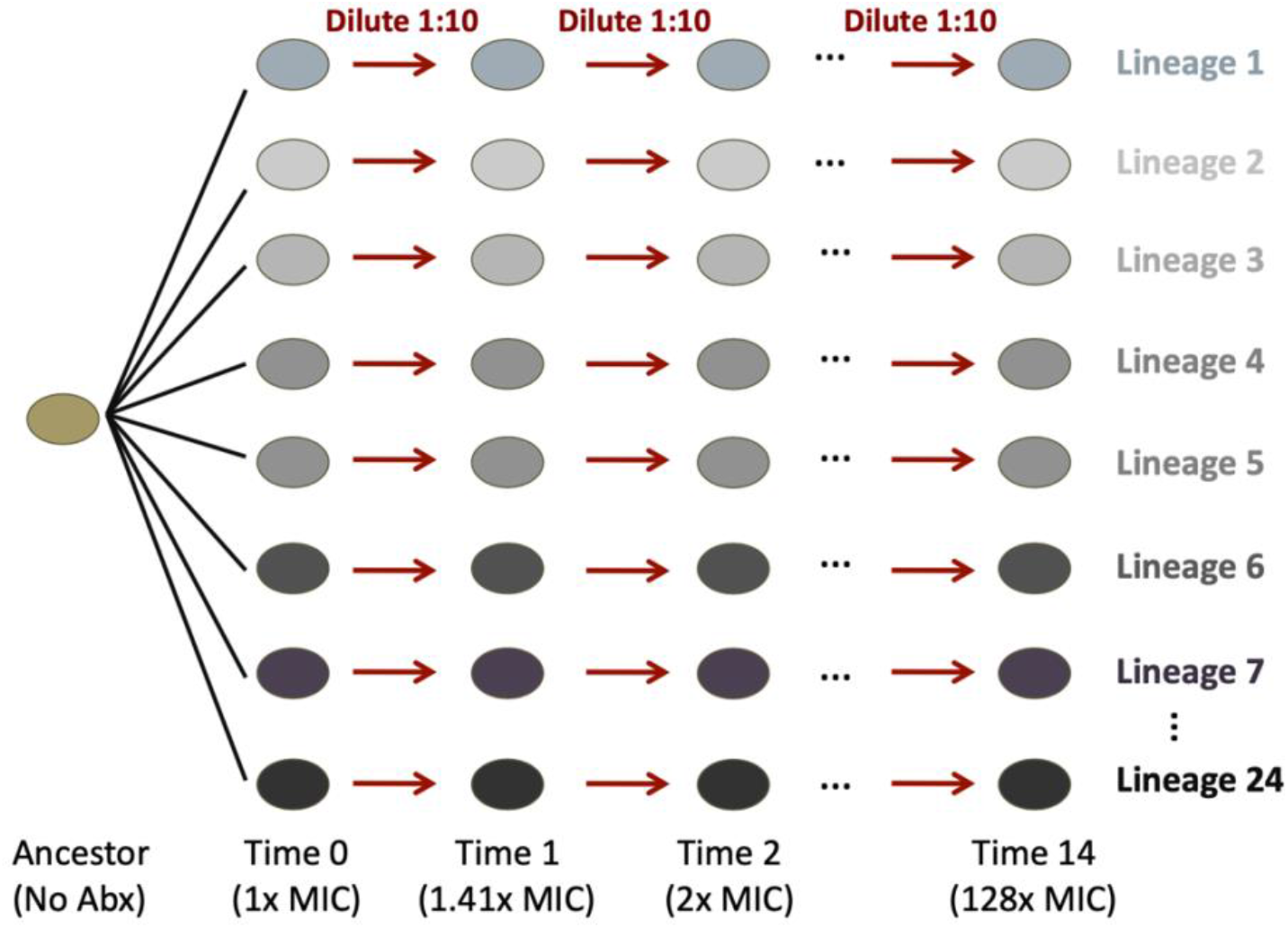
Schematic of evolution experiment to obtain strains of varying chloramphenicol resistance. Beginning with a common ancestor, we first passaged cultures at their MIC to create 24 replicate lineages. When transferring cultures to new media, we diluted cultures 1:10 during each passage. The rate of increase in antibiotic was a factor of √2 between sequential levels.

### Resistance incurs costs in both thermal tolerance and maximum growth rate

We measured growth rates of experimentally evolved *E. coli* cultures at three different temperatures: their historic temperature of 37C, and the novel temperatures of 32C and 42C. We hypothesized that growth rate costs of resistance would be larger in the novel temperatures, consistent with reduced thermal niche breadth.

Overall, we found the growth rates decreased strongly with increasing antibiotic resistance (Fig. 3, left). We then calculated relative growth rates for each lineage by dividing the growth rate at each timepoint by the growth rate of the culture at timepoint 1 (T1). Analysis of these relative growth rates showed that there was both a fitness cost in maximum growth rate and a fitness cost in thermal niche breadth; the linear model showed a strong negative effect of increasing resistance on growth rate at 37C (F1, 974 = 988.2, p < 0.001), and significant variability in the effect of resistance on growth at the three different temperatures (F1, 974 = 13.8, p < 0.001). The negative effect of resistance on growth rate was greater at 32C and 42C than at 37C (Fig. 3, right). Thus, there were disproportionate fitness costs in both the novel temperatures. This finding is corroborated by counting the number of evolved populations that increased their growth rate, relative to the ancestor; at 37C, there were 41 cultures with increased growth rates, whereas at 32C there were 13 and at 42C only 2. Therefore, the presence of fitness costs was more consistent in the novel temperatures.

**Figure 3:**
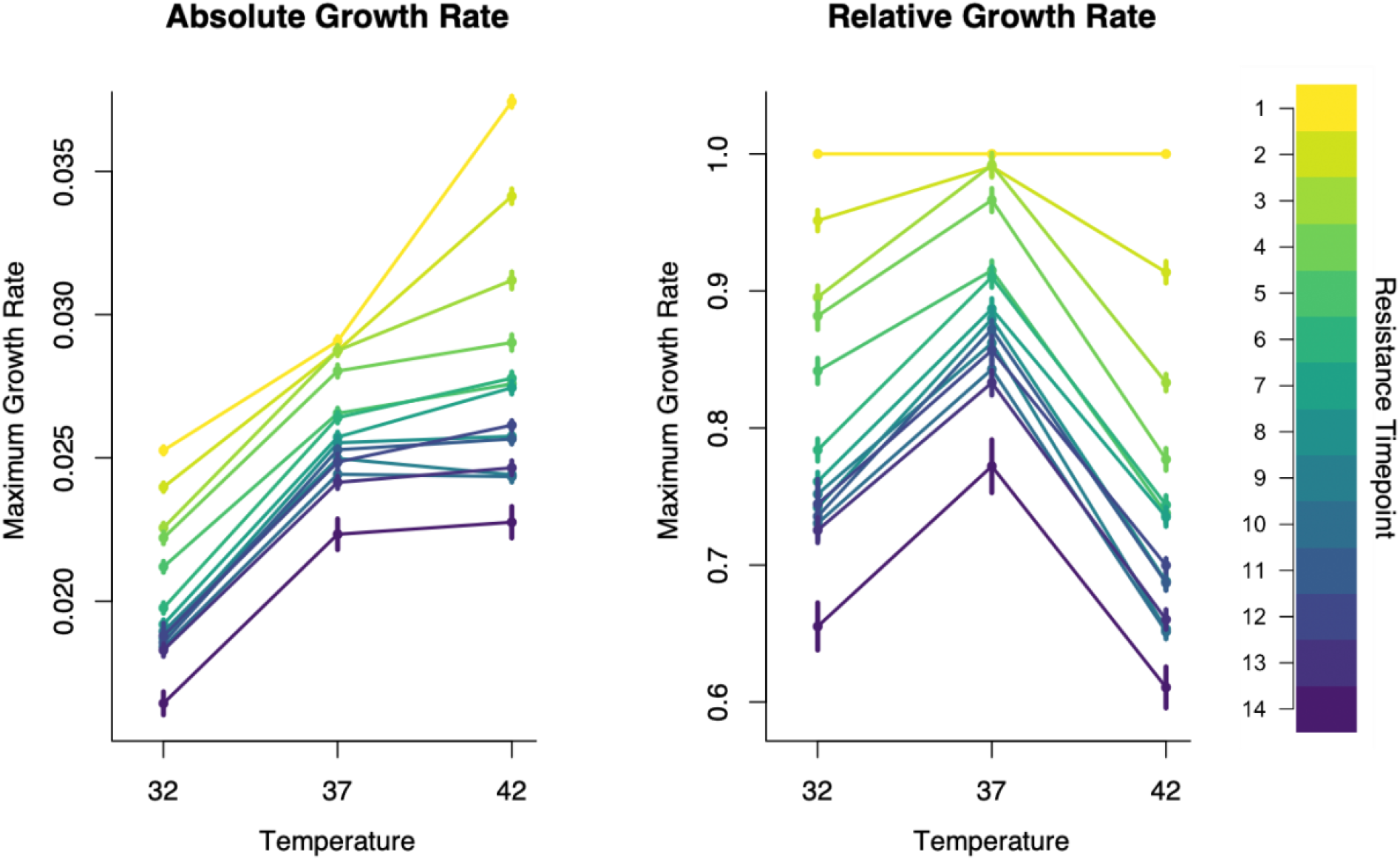
Strong fitness costs to both thermal niche breadth and maximum growth rate. Absolute growth rates varied across the three temperatures, and were generally highest at 42C for a given culture (left). Increasing resistance, as measured by timepoint in the evolution experiment (darker colors indicate higher resistance levels), resulted in reduced maximum growth rates (left). These differences are more apparent when analyzing relative growth rates (i.e. the growth rates divided by the respective ancestral growth at T1). As resistance increased, the relative growth rate decreased at 37C, but showed even larger declines at the novel temperatures of 32C and 42C. Of the three scenarios depicted in Fig. 1, these findings mirror the right-hand panel in Fig. 1, where there are costs to both maximum growth rate and thermal niche breadth. Points show the mean and standard error of growth rates from measurements from each of the 24 lineages.

### Strong competitive disadvantage of resistance at increased temperature

Next, we evaluated the effects of increasing chloramphenicol resistance on competitive success, measured by the fraction of the community comprised by a resistant strain when grown in mixed culture with a more sensitive strain. We transformed strains from 96 cultures (timepoints 1, 5, 9, and 13 for each of the 24 lineages) with either GFP or mCherry plasmids and quantified population sizes after competition assays using flow cytometry. We found that resistance level was a strong driver of the composition of mixed cultures in the competition experiments (Fig. 4). For example, at 32C and 37C, the most sensitive strains (T1) grew to a population 1.3x larger when grown against the most resistant strain (T13), as compared to being grown against the same timepoint (T1). The growth differential was much stronger at 42C, where the most sensitive strain (T1) grew to 3x their population size when competed against the most resistant strain (T13).

**Figure 4:**
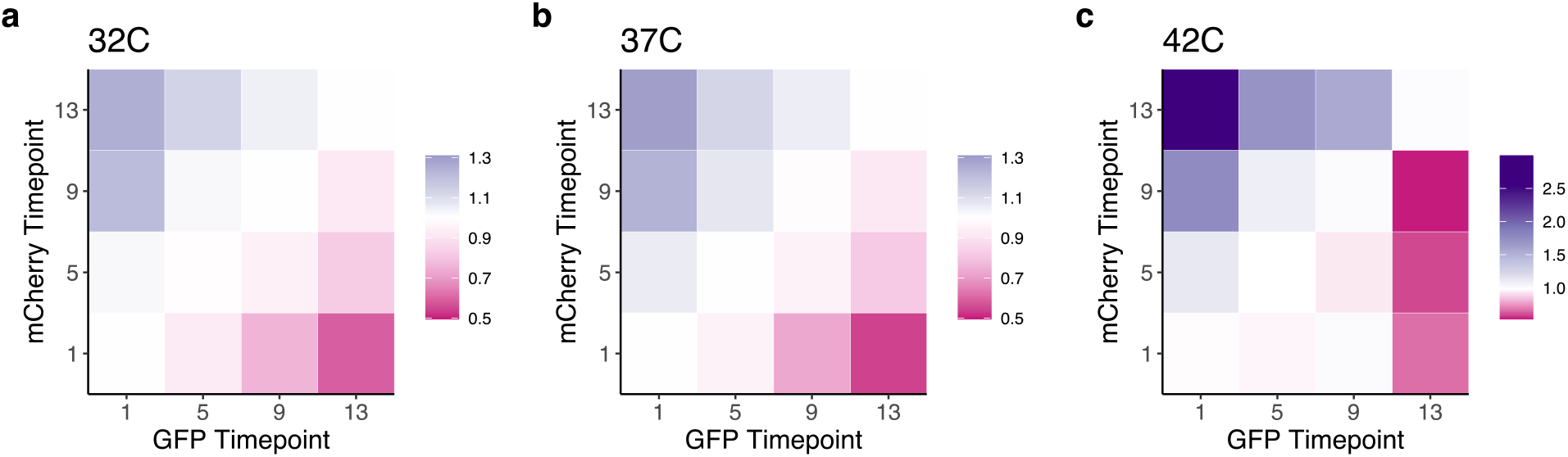
Resistant strains are outcompeted by their sensitive ancestors in the absence of drug, particularly at the warmer temperature. Heatmaps show the performance of strains in competition experiments at the 3 temperatures (ratio of its performance in competition with another strain versus competition with itself), where the x-axis is the timepoint of the GFP strain and the y-axis is the timepoint of the mCherry strain. Purple cells indicate that the GFP strain out-performed in final community composition, as compared to when that same GFP strain was competed against a strain from the same timepoint, but with the mCherry plasmid. In contrast, darker pink cells indicate that the GFP strain performed worse than in competition with the same timepoint. At 42C (panel c), the more evolved strains had stronger differential success than at 32C or 37C (a and b), indicated by the darker colors (note the different scale bars between 42C and the other two temperatures).

For each lineage, we evaluated whether the competitive cost of resistance was different at 32C or 42C when compared to the effect at 37C. We found that 12 of the 24 lineages had significantly stronger negative effects of resistance on strain growth at 42C, as compared to 37C. One lineage (L16) had significantly smaller costs of resistance at 42C. There were no lineages where there was a significant difference in the effect of resistance between 37C and 32C. Full results can be found in Table S1. When removing the interaction between temperature and lineage to evaluate the average effect of temperature across all lineages, we found that the effect of resistance differed across temperatures (F2, 3309 = 51.3, p < 0.001). Specifically, the negative effect was greater at 42C than 37C (a slope 59% greater at 42C), though there was no significant difference between 32C and 37C (a slope 11% weaker at 32C). To visualize these differences in the effect of resistance on strain growth, we show two lineages with contrasting results; lineage 5 had no significant differences in the effect of resistance across temperature, whereas lineage 19 showed a much stronger effect of resistance differences at 42C than at the other two temperatures (Fig. 5).

**Figure 5:**
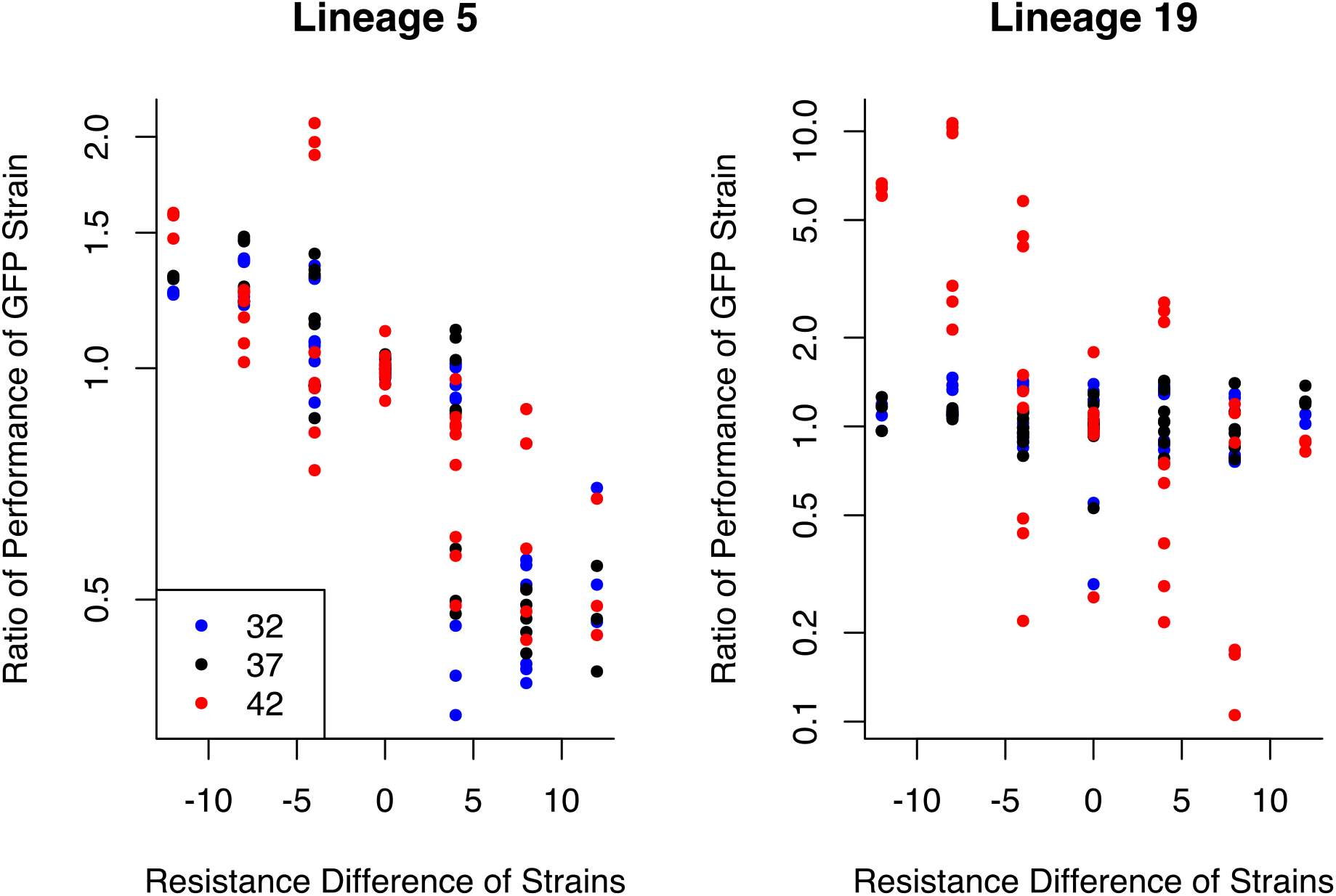
Lineage-specific fitness costs in thermal tolerance. While many lineages demonstrated a greater cost of resistance at 42C than the other two temperatures, some lineages showed no increased effect at the warmer temperature. Lineage 5 (left) is an example of an evolutionary trajectory where fitness costs were not substantially different across temperatures. The x-axis is the difference in experimental timepoints at which strains were collected (a proxy for minimum resistance level). The y-axis gives the change in performance of the GFP strain as compared to competition against the same timepoint. Conversely, lineage 19 (right) shows an example of a strong interactive effect of fitness costs and temperature, where strain performance is more impacted by resistance at 42C (red circles) than at the other two temperatures.

### Variability in routes to resistance among lineages

Finally, we sequenced genomes for the 96 strains used in the competition experiments to evaluate how the genetic mechanism of resistance affected fitness. Using the genomic data, we identified 220 mutations across these 96 strains; of these mutations, 24 occurred within strains at T1, 44 in strains at T5, 65 in strains at T9, and 87 in strains at T13. We also identified the timepoint at which these mutations were first identified within a lineage. All 24 mutations at T1 were new when comparing against the ancestor, along with 29 mutations present at T5 but not the same lineages at T1, 26 mutations present at T9 but not T5, and 40 mutations present at T13 but not T9. We saw that most mutations fell within genes for known resistance mechanisms, such as efflux pumps, the multiple antibiotic resistance protein (“mar”), or ion channels. We grouped the mutations into 10 categories for further analysis: those associated with the acr efflux pump (64 mutations), ATP synthase (42 mutations), the mar resistance protein (20 mutations), the mdfA efflux pump (15 mutations), other metabolism (generally related to carbon usage, 16 mutations), the mscK ion channel (12 mutations), outer-membrane-associated proteins (12 mutations), mutations in prophages (15 mutations), slt degragation protein (12 mutations), and mutations affecting transcription/translation (12 mutations). We analyzed the dataset containing the first observation of each mutation to assess whether certain mechanisms appeared earlier or later in evolutionary trajectories (Fig. 6). Using chi-squared tests on the number of mutations in each category across the four timepoints, we found that mutations associated with the acr efflux pump (p < 0.001), ATP synthase (p = 0.041), the mar resistance protein (p = 0.022), outer membrane proteins (p = 0.018), and the mfdA efflux pump (p = 0.013) were nonrandomly distributed across evolutionary time. Specifically, early mutations at T1 were disproportionately located within the acr efflux pump, while mutations at T5 were often within the acr efflux pump and associated with ATP synthase. At T9, mutations within the outer membrane proteins became more common. Finally, the most resistant strains frequently had mutations in the mar resistance protein and the mfdA efflux pump. Although conducting 10 chi-squared tests for the 10 categories allows the possibility of finding spurious positive results, the probability of spuriously identifying five categories at a threshold of p < 0.05 is less than 1 in 16,000.

**Figure 6:**
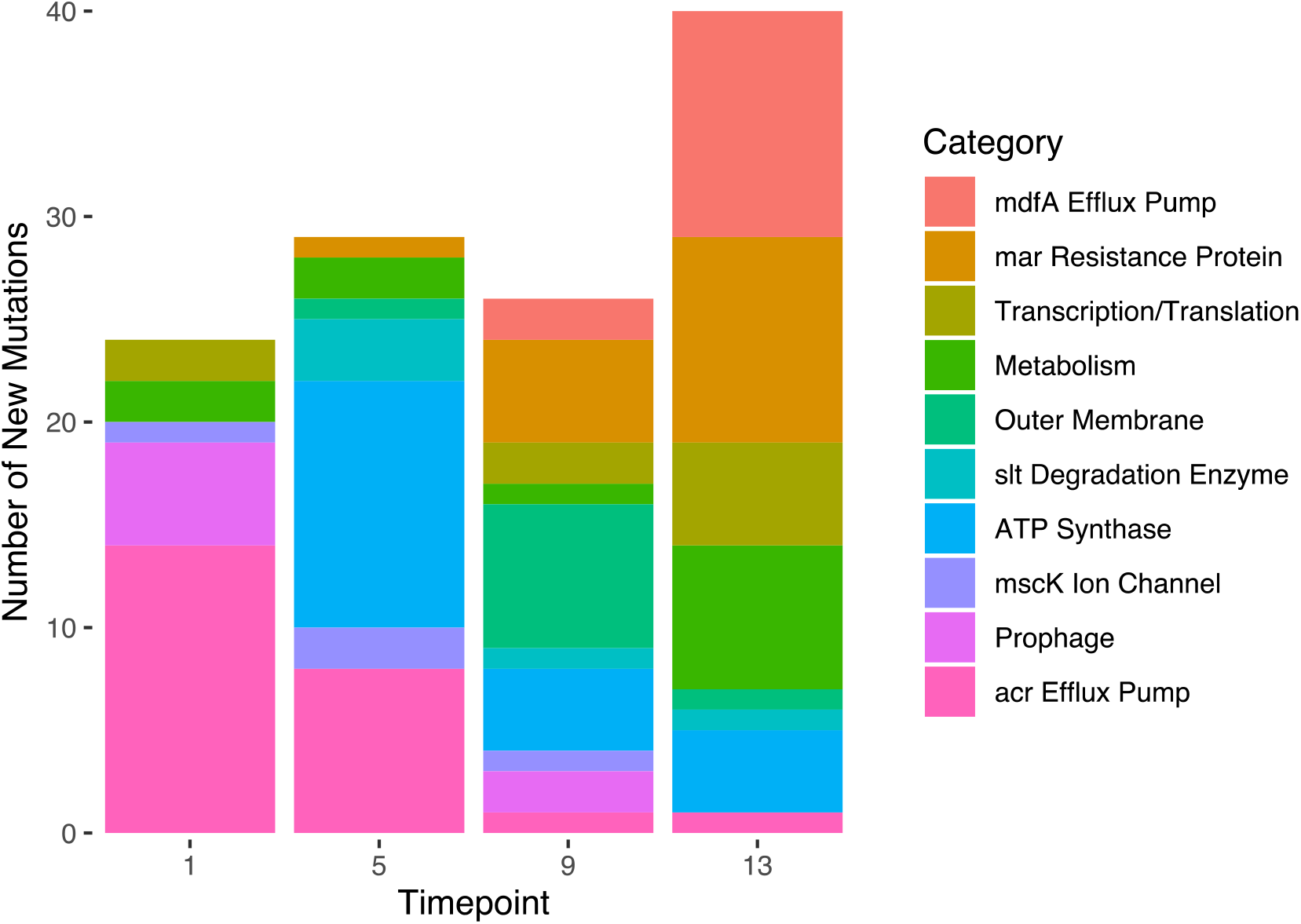
Transition of resistance strategies across the chloramphenicol gradient. The types of mutations acquired at each timesteps changed as antibiotic levels increased. The most common pathways to resistance at the four timepoints were mutations in the acr efflux pump (T1), ATP synthase genes (T5), outer membrane proteins (T9), and the mdfA efflux pump (T13).

### Fitness effects of resistance mutations are more extreme at higher temperature

We calculated the estimated effect of mutations in 37 genes on the outcomes of the pairwise competition experiments, as to compare the distribution of mutational fitness effects across the three temperatures. Many of these genes were part of the same cassettes (e.g. acrA, acrB, and acrR). Of these 37 genes, we saw significant differences in the estimated effects of mutations in 24 genes between 37C and 42C. Conversely, there were zero significant differences in effect size between the temperatures of 32C and 37C. Effect sizes in this analysis signify the difference in the success of a strain as a result of carrying one extra mutation in the indicated gene. Then, we evaluated whether the range of gene effects was larger at 42C or 32C, as compared to the range of gene effects at 37C (Fig. 7). We conducted F-tests on the distribution of gene effect sizes at 37C versus 42C and at 37C versus 32C to find whether the variances of these distributions were unequal. We found that the range of gene effects was greater at 42C than 37C (F36, 36 = 5.68, p < 0.001) but that the range of gene effects was not different between 32C and 37C (F36, 36 = 0.91, p = 0.79).

**Figure 7:**
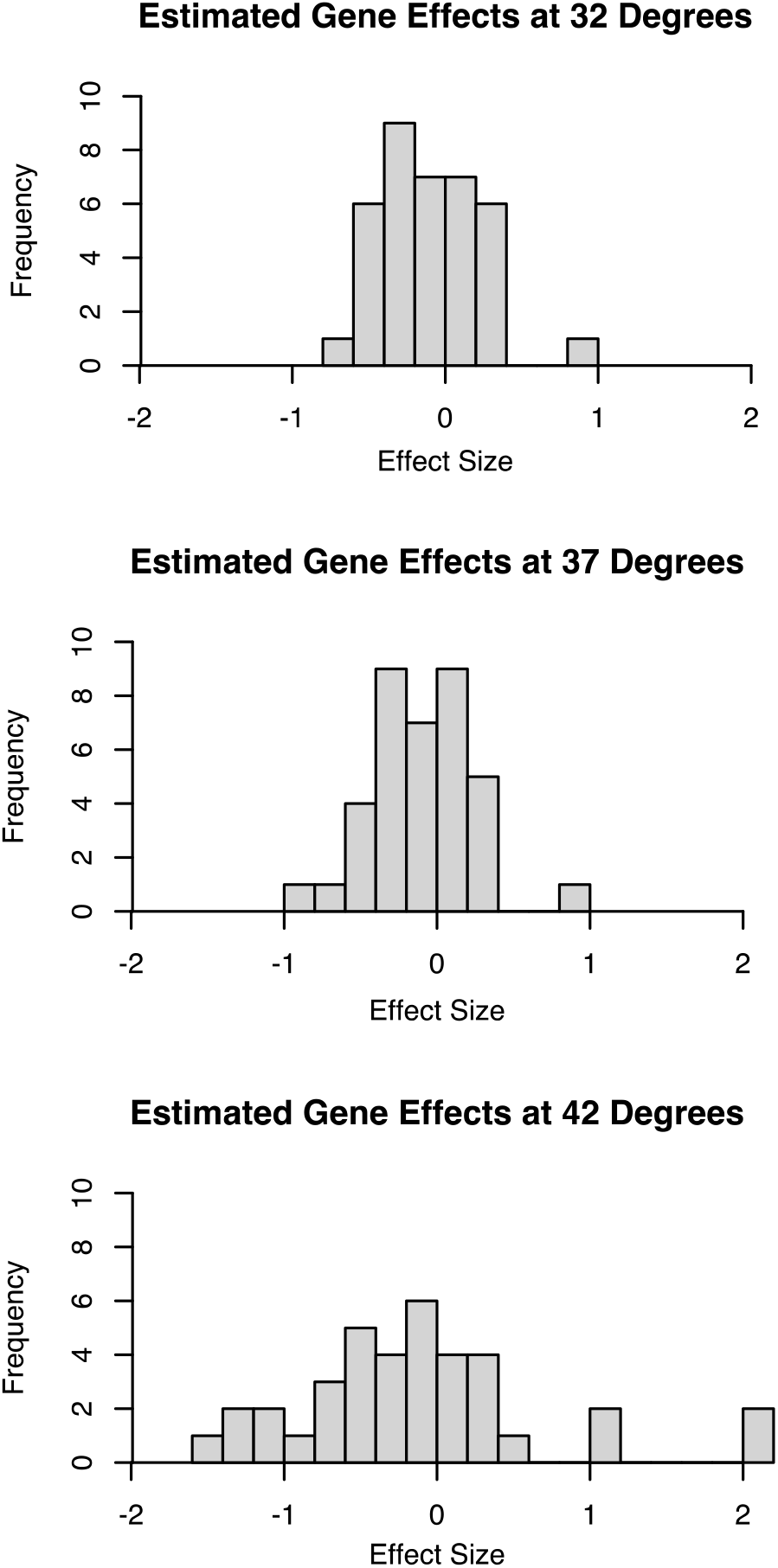
Effects of mutations are more extreme at a warmer temperature. Results of the linear regression quantifying the effects of mutations within 37 genes show that the estimated impact of mutations is greatest at 42C, as compared to 32C or 37C. Effect sizes near zero indicate no effect, whereas an effect size of 1 indicates a one-log-fold increase in the success of a strain that has acquired a mutation in the given gene. The variance of effect sizes at 42C is significantly greater than at the other two temperatures.

The genes with the most significant additional fitness cost at 42C were: mscK, opgH, dgcF, atpD, uvrA/ssb, and ssb. Of these, only mutations in atpD and mscK were common, appearing in more than 1 lineage. These genes are involved in ATP synthesis and in ion transport via the mechanosensitive channel MscK, respectively.

Finally, we combined the competition data with the genomic data to examine whether specific categories of mutations were more likely to result in decreased thermal tolerance. For this analysis, we used Fisher’s Exact Tests to determine whether lineages with a greater cost of resistance at 42C (12 lineages) were more likely to have any specific type of mutation (using the 10 mutation categories shown in Fig. 5). Surprisingly, of these 10 tests, we found no significant (p < 0.05) associations between mutation type and the presence of fitness costs to thermal niche breadth, suggesting that the tradeoff is not tied to a specific mechanism of resistance.

## Discussion

Taken together, the results of our experiments show strong evidence for a narrowing of thermal tolerance as a trade-off of antibiotic resistance. Specifically, fitness costs in growth rate were greater in novel temperatures (32C and 42C) than at the historic temperature of 37C, indicating that resistant bacteria are less able to tolerate these new environments (Fig. 3, Fig. 4). Trade-offs to antibiotic resistance are particularly strong at 42C, where population dynamics appear qualitatively different than 37C and 32C. At 42C, costs in growth rate were nearly ubiquitous among lineages, and resistant strains composed a much smaller fraction of community composition in the competition experiments. Interestingly, no single genetic strategy of resistance seemed more likely to cause this decreased thermal tolerance, suggesting that this fitness cost might be generalizable to other antibiotics with different resistance mechanisms. Although the growth rate and competition experiments were largely concordant, there were minor differences in the effects of lower temperature on apparent fitness costs, which may be explained by several factors. First, there are possibly multiple coexisting strains in growth rate cultures, but only one genotype was selected for the competition experiments. Noting the large spread in the effect of resistance mutations on fitness (Fig. 7), the outcome of competition experiments depends on the specific genotypes, in addition to the resistance level. Second, as noted previously, additional factors such as resource use efficiency and resource uptake rates also affect eventual population size. Overall, our experiments demonstrate that chloramphenicol resistance reduces the ability of evolved cells to grow in novel thermal environments, and that these trade-offs put resistant strains at a strong competitive disadvantage at warmer temperatures.

Despite the strong overall pattern that resistant strains fared poorly at higher temperatures, there was substantial variation between lineages in this effect. There are many variables that could mediate the fitness costs across lineages, though the most obvious is that the stochastic accumulation of mutations leads to varied genetic bases of resistance with varied costs in new environments (Kassen and Bataillon 2006). Thus, future experiments studying the trade-offs in novel environments should include a high degree of replication in order to determine the probability of such costs existing in a single lineage. There are additional experimental factors that could interact with this stochasticity of evolutionary trajectories, including population size and the strength of the selection gradient. At very large population sizes, convergence of evolutionary trajectories becomes more likely due to mutation saturation within populations (Szendro et al. 2013); in this case, the most strongly beneficial mutations arise in each population, and will persist. We can be sure that these experiments did not approach mutation saturation because only one lineage evolved the strongly beneficial media adaptation in araD. This mutation can be clearly seen as a positive outlier in the estimated gene effect sizes at 37C, yet it only appeared once (Fig. 7). Similarly, the strength of the selection gradient determines the fraction of the population that can survive passaging to a higher antibiotic concentration. At stronger selection gradients, fewer genotypes would survive the incremental increase in antibiotic, leading to a smaller number of genetic routes to resistance (Wistrand-Yuen et al. 2018). Furthermore, the specific media and antibiotic used also influence the distribution of fitness costs, as the variety of carbon sources and the specific antibiotic targets also determine the number and type of mutations conferring fitness advantages. Finally, it would also be interesting to evaluate whether structural variation in genomes (such as duplications or inversions) or changes to transcriptional regulation correspond to the observed fitness costs, as our present sequencing methodologies cannot address these possibilities.

Two components of our analyses suggest potential mechanisms leading to decreased thermal niche breadth: the mutations involved in the genetic basis of resistance, and the result that warmer temperatures are more detrimental to resistant bacteria. Increased temperatures have a multitude of effects on bacterial physiology, including increased membrane fluidity (de Mendoza and Cronan 1983), altered protein folding (Chowdhury et al. 2006), reduced generation time (Ratkowsky et al. 1981), and increased speed of chemical reactions (Ratkowsky et al. 2005). These effects of temperature might interact with specific mechanisms of resistance to yield the disproportionate fitness costs. For example, many proteins involved in resistance are embedded within the outer membrane, such as efflux pumps and outer membrane porins. The efficacy of these resistance mechanisms, or the energy needed for these complexes to function, might change as a function of membrane fluidity. A higher temperature could also increase cell susceptibility to antibiotics, as altered protein folding at this temperature might expose additional chloramphenicol targets within ribosomes. In this case, cells might need to upregulate activity of resistance mechanisms to produce the same level of resistance. Alternatively, the location of mutations within proteins might be more detrimental at 42C due to altered protein structure. Furthermore, the ATP synthase mutations that were observed may confer resistance by slowing cellular metabolism sufficiently that chloramphenicol targets are no longer active; this is frequently seen in persister cells (Lewis 2005), and persister mutations are common in early stages of resistance evolution (Windels et al. 2019). It is possible that there is a greater cost of these ATP synthase mutations at 42C, as higher temperatures often require increased cellular metabolism and respiration. Finally, the rapid drop-off in activity of many enzymes above their thermal optimum (Elias et al. 2014) means that cells may be particularly sensitive to perturbations near the upper bound of their thermal range. Thus, a resistance mutation that changes the enzyme kinetics by a small amount would have the greatest effect near this upper thermal limit.

In addition to thermal tolerance, there are many other dimensions of niche breadth where fitness costs could manifest. Further exploration of environmental tolerance might include the ability to withstand desiccation or changes in pH. Another aspect of decreased niche breadth could be the ability to survive biotic interactions, such as the ability to evade a wide variety of immune system responses or to avoid infections by bacteriophages. Of particular importance are trade-offs that would manifest in reduced virulence. This category of fitness costs to niche breadth might include reduced host range or decreased survival outside the human body. These two dimensions of fitness would slow the spread of person-to-person infection by either decreasing the number of susceptible hosts or decreasing the transmission rate of an infected individual. Thus, further studies of evolutionary fitness costs might explicitly incorporate multiple stages of a pathogen’s life cycle. More broadly, the need to simultaneously maintain adaptations under antibiotic pressure reflects the pressures of a generalist versus specialist strategy. Theory indicates that it might be difficult to evolve strong specialization (e.g. antibiotic resistance) without losing other functionality due to pleiotropic effects (Bono et al. 2019). Because the long-term survival of resistant populations is the multiplicative effect of survival across different life stages, modest trade-offs in niche breadth could strongly impact the population dynamics of resistant bacteria.

Given the impact of costs to niche breadth on the frequency of resistant bacteria, these trade-offs likely have consequences for the long-term evolution of antibiotic resistance. For example, if narrower niche breadth means that resistance can change from beneficial to detrimental over short time spans (as is the case when changing environments), this rapid reversal in fitness could select for easy gain and loss of resistance. Indeed, this is seen frequently in bacteria, when resistance in nature can be nearly instantaneously acquired by horizontal gene transfer or transposable elements (Partridge et al. 2018). Additionally, these experiments suggest that there are further dimensions of bacterial fitness that have not been tested by laboratory experiments, due to the homogenous environments used for cell culturing. Testing for fitness costs in novel environments could shed light on the temporal or spatial distribution of resistance evolution. For example, the finding that resistance decreases thermal niche breadth might contribute to the observation that resistance increases with ambient temperature (MacFadden et al. 2018). Consequently, these experiments raise the question of how adaptation for resistance interacts with adaptation to other specific stressors. For example, it would be both clinically and theoretically valuable to know whether antibiotic resistance is lost more readily when cells are subjected to novel selective pressures. More broadly, integrating niche breadth into the framework of fitness costs has the potential to resolve seeming contradictions in resistance evolution; costs in niche breadth allow for the possibility that any resistance mutation has trade-offs, but the costs may not be apparent in a constant environment.

## Methods

### Obtaining resistant strains via experimental evolution

Protocol: Beginning with a single liquid culture of Keio strain ΔlacA (Baba et al. 2006), we first identified the MIC of this strain (~3ug/mL) by testing its growth in M9 with varying concentrations of chloramphenicol. We passaged these cultures at a concentration of 2.5ug/mL to allow populations to diversify before beginning the selection experiment. We initiated 96 cultures in a 96-well plate, and selected the 24 wells with strongest cell growth to propagate further. Experiments were done using M9 minimal media to preclude biphasic growth arising from multiple potential carbon sources. The cultures were incubated at 37C with continuous shaking. For each passaging step, the following protocol was used:

1. Ensure that all 24 cultures have visible growth
2. Passage one-tenth of the culture to new media containing a chloramphenicol concentration of ✓2x of the prior concentration
3. Allow cultures to grow for 48h
4. If there is not visible growth in all cultures, passage the cultures at the same chloramphenicol concentration and allow cultures to grow for 48h

Cultures were initiated in 96-well plates with 150ul volume, but were moved to 5mL cultures at timepoint 6, because the small populations were easily extinguished with increasing antibiotic concentrations. As chloramphenicol concentrations became very high (>50x initial MIC), transfers to higher antibiotic concentrations strongly inhibited growth. In the case that there was no visible growth after passaging at the same chloramphenicol concentration, we passaged all cultures in media with no chloramphenicol to restore cell density. Then, we resumed the typical protocol steps, beginning at the concentration the cells previously experienced. We terminated the experiment when cultures were no longer able to grow when transferred to a higher chloramphenicol concentration, which occurred after 14 timepoints, representing 128x the starting concentration.

Cultures were collected for archival storage immediately before being passaged to media with a greater chloramphenicol concentration. We restored cell densities by growing each culture for 48h in the absence of antibiotic, and stored resulting cultures at −80C in 20% glycerol.

### Quantifying fitness: measuring maximal growth rates

We quantified bacterial growth rates by measuring the turbidity of cultures (OD 600) using a BioTek Synergy H1 plate reader. We measured 60 cultures per 96-well plate, using only the interior wells of the plate. We used a 96-well pipettor to inoculate 150ul of fresh media with 2ul of stored culture. Cultures were incubated at a constant temperature for 24 hours with continuous shaking, and measurements taken every 5 minutes. The growth of each culture was measured at three temperatures: their historic temperature of 37C, and the novel temperatures of 32C and 42C.

The measurement of interest in these experiments was the maximum growth rate achieved by each culture at each of the three temperatures. To process and clean the raw data, we first re-centered the optical density data to have a minimum value of 0.02, indicating a small population at the outset of the measurements. This was necessary because the media and the plate lid caused optical density measurements of approximately 0.1, even in wells with no bacterial cells. Results of the subsequent analyses were robust to changes in the value selected for the initial optical density. Then, we used a smoothing function to minimize the effect of discontinuities in optical density measurements, which were generally small but could have disproportionate effects on the data at low optical density values; we used a local linear model using the surrounding 24 data points to smooth the values. Optical density values were log-transformed before smoothing. Again, results were robust to varying the number of data points included to produce the smoothing. Finally, to find the maximum growth rate for each sample, we subtracted sequential values after log-transforming and smoothing, and used the maximum difference as the measurement of the maximum growth rate.

We hypothesized that there would be both costs in maximum growth rate and costs in thermal niche breadth as bacteria became more resistant to chloramphenicol. We used a linear regression to evaluate this prediction, where the outcome variables were the growth rates of the cultures (24 lineages x 14 timepoints x 3 temperatures = 1008 growth rate measurements) and the predictors were the temperature of measurement (32C, 37C, or 42C), lineage (L1 - L24), the timepoint (1 through 14, representing increasing chloramphenicol resistance), and an interaction term between temperature and timepoint. This interaction tests whether the effect of timepoint on growth rate is differentially strong at the three temperatures. In the five instances where there was minimal growth in the cultures (change in OD600 < 0.05), we removed the data points from the analysis.

### Competing evolved versus ancestral strains

Next, we wanted to directly evaluate how increasing chloramphenicol resistance influenced competitive ability between ancestral and evolved bacterial strains. This question addresses how the acquisition of antibiotic resistance influences the frequency of the resistant strain in mixed populations. We competed genotypes from the same lineage but different timepoints against each other in the same well of a 96-well plate. The timepoints used for competition experiments were T1, T5, T9, and T13, with all competition experiments done in a pairwise fashion and replicated three times.

We quantified the population sizes of the different strains at the various timepoints via flow cytometry. For each strain used in the competition experiments (24 lineages x 4 timepoints = 96 strains), we created two transformed strains with either the pFCcGi plasmid to express the mCherry protein or the pDiGc plasmid to express the GFP protein. These plasmids confer resistance to ampicillin, and are suited for competition experiments because they are similar plasmids that were constructed by the same research group (Helaine et al. 2010, Figueira et al. 2013). We selected one colony from the successful transformations, and created new archived stocks of these strains by growing these strains for 48h in M9 media containing ampicillin and then storing them at −80C in 20% glycerol.

For the competition experiments, wells in a 96-well plate were filled with M9 containing ampicillin and were inoculated with 2ul of two strains. In each well, there was one strain with the GFP-producing plasmid, and one strain with the mCherry-producing plasmid. Competition experiments were conducted at 3 temperatures (32C, 37C, and 42C) by incubating the 96-well plate at the given temperature for 24h with continuous shaking. At 24h, plates were removed from the incubator, and 6ul of each mixed culture was transferred to 200ul of water to be read on an LSRII flow cytometer. A minimum of 10,000 cells were counted for each well. The outcome of interest was the population success of the GFP-producing strain; we analyzed the GFP-producing strains, because the population changes were more symmetrically distributed around zero with less skewness. These experiments produced 3,456 measurements of population size (4 GFP strains x 4 mCherry strains x 3 replicates x 3 temperatures x 24 lineages).

Our hypothesis for the competition experiments was that the more resistant strains would fare worse at novel temperatures (32C or 42C) than in their historical temperature (37C). We evaluated this by comparing the fraction of the community comprised of each GFP-transformed strain as a function of the difference in resistance (measured by the difference in the experimental timepoints of the two competing strains). We conducted a linear mixed-effects regression where the outcome variable was the log-ratio of the community comprised by the GFP strain divided by the average fraction of the community comprised by the GFP strain when grown together with the same timepoint with the mCherry plasmid. This standardization accounts for potential differential costs of carrying the GFP versus mCherry plasmids. A value of 1.4 for the well where lineage 3 T5 containing GFP was competed against lineage 3 T13 mCherry would indicate that the lineage 5 T5 GFP strain comprised 1.4 times the fraction of the community than it did when competed against lineage 3 T5 mCherry strain. The fixed effect variables in the regression included the difference in resistance between strains, temperature, and lineage, with all interaction terms included. The difference in resistance was measured by the difference in the experimental timepoints of the two competing strains. For example, if the GFP-transformed strain were from T5 and the mCherry-transformed strain were from T13, the difference in resistance would be −8. A significant interaction between temperature and resistance difference indicates that the effect of resistance on strain growth is different among temperatures. We included a random effect to allow the log-ratio of GFP strains to vary in response to competing against different mCherry strains.

### Genomic sequencing to identify genetic basis of resistance

After selecting strains for the competition experiments, we sequenced the GFP-transformed versions of these 96 strains. We followed the protocol set out in Baym et. al 2015; briefly, we extracted genomic DNA using an Invitrogen kit, then carried out tagmentation using the Illumina Nextera kit. We used Q5 High Fidelity 2x Master Mix for the library prep, and cleaned up the libraries with SPRI magnetic beads. We ran the sequencing on a Novaseq at the Harvard Bauer Core, producing 150bp paired-end reads. The minimum number of reads for a given sample was 2.4million, resulting in coverage of over 100x for all samples. We also sequenced the initial ancestral strain prior to exposure to any antibiotic.

We used the computational pipeline, breseq, to determine mutations arising in each strain (Deatherage and Barrick, 2014). This toolkit was developed for the purpose of analyzing microbial genomes during the course of evolutionary experiments. We used a genome for *E. coli* K-12, substr. MG1655 (#U00096 from Genbank) as the initial reference, which we updated using the data from our sequenced ancestral strain. The output of this pipeline identifies locations of mutations in each genome and specifies the genes in which they occur.

Finally, to examine whether there were differential effects of mutations across the three temperatures, we analyzed whether the location of mutations within each strain could explain the outcome of the competition experiments described above. We used a linear regression where the outcome variable was, again, the log-ratio of the strain in competition divided by the population of the strain when grown against the same timepoint. The predictor variables were the difference between the two strains in the number of mutations in each gene (e.g. L11 T13 has 1 mutation in marR and L11 T9 has zero mutations in marR, the value for marR for L11 T13 is 1). We included interactions between all genes and temperature, in order to test whether estimated effect sizes of mutations differed in the three thermal environments. Also, we included lineage as a predictor variable to account for potential differences in the success of the GFP strains between lineages. We identified mutations in 43 genes, though the distribution of mutations in 6 of these genes (yejA, yfaQ, xdhB, rplD, rpoC, and uvrA) were equivalent to the mutation occurrence of other genes, and thus could not be included in the analyses due to the fact that the predictor variables were identical.

## Supporting information

Merged Flow Cytometry Data

Growth Curve Data

Annotated Mutation Data

Document with R scripts for analysis

## Acknowledgments

We thank Anurag Limdi, Eleanor Rand, and Karel Brinda for their constructive comments. Flow cytometry analyses were conducted at the Harvard Medical School Systems Biology FACS facility. This work was supported by the NIH NIGMS award R35GM133700, the David and Lucile Packard Foundation, the Pew Charitable Trusts, and the Alfred P. Sloan Foundation. CH was partially supported by the Harvard Data Science Initiative.

