## Supplementary material for "Decreased thermal tolerance as a trade-off of antibiotic resistance": Document with R scripts for analysis

```
## Analyze merged flow cytometry data

library(lme4)
library(RColorBrewer)
library(ggplot2)
library(dplyr)
library(gridExtra)
library(nlme)

all.ev <- read.csv("~/Desktop/MergedFlowCytometryDataset.csv")

# Relevel temperatures so linear model compares against 37 as baseline
all.ev$temp <- as.factor(all.ev$temp)
all.ev$temp <- relevel(all.ev$temp, ref = "37")

# Conduct analyses of log-ratios
summary(lme(log(gfp.rawratio) ~ -1 + lineage * gfp.timediff * temp , random =  ~ 1|mchstrain, data = all.ev) )
## Count which lineages have interaction:
## 1, 2, 7, 10, 11, 13, 15, 18, 19, 20, 21, 22

# Collapse all temperature interactions by lineage to see overall effect of temperature
summary(lme(log(gfp.rawratio) ~ lineage * temp + gfp.timediff * temp , random =  ~ 1|mchstrain, data = all.ev) )
anova(lme(log(gfp.rawratio) ~ lineage * temp + gfp.timediff * temp , random =  ~ 1|mchstrain, data = all.ev) )

#Make plots of same lineage at different temperatures

par(mfrow = c(1, 2))
lineage.plot1 <- "L5"
plot(all.ev$gfp.rawratio[all.ev$lineage == lineage.plot1] ~ all.ev$gfp.timediff[ all.ev$lineage == lineage.plot1], xlab = "Resistance Difference of Strains", ylab = "Ratio of Performance of GFP Strain", log = "y", pch = 20, main = "Lineage 5", col = rep(c("blue", "black", "red"), each = 48) )
legend("bottomleft", bty = "l", legend = c("32", "37", "42"), col = c("blue", "black", "red"), pch = 20)

lineage.plot2 <- "L19"
plot(all.ev$gfp.rawratio[all.ev$lineage == lineage.plot2] ~ all.ev$gfp.timediff[ all.ev$lineage == lineage.plot2], xlab = "Resistance Difference of Strains", ylab = "Ratio of Performance of GFP Strain", log = "y", pch = 20, main = "Lineage 19",  col = rep(c("blue", "black", "red"), each = 48) )

## Next, make heatmaps of gfp ratios at the 3 temperatures. Should be 4x4 grid of GFP and mCh concentrations

# First split up tapply names to get resistance levels
all.ev$combo_temp <- paste(all.ev$gfptime, all.ev$mchtime, all.ev$temp, sep = "_")

name.split <- as.data.frame(strsplit(names(tapply(all.ev$gfp.rawratio, all.ev$combo_temp, mean)), "_"))
name.split <- rbind(name.split, tapply(all.ev$gfp.rawratio, all.ev$combo_temp, median) )
name.split <- t(name.split)
colnames(name.split) <- c("gfp.time", "mch.time", "temp", "ratio")
name.split <- as.data.frame.matrix(name.split)
name.split$log.ratio <- log(as.numeric(name.split$ratio))

name.split32 <- name.split[name.split$temp == "32", ]
name.split37 <- name.split[name.split$temp == "37", ]
name.split42 <- name.split[name.split$temp == "42", ]

gfp.levels <- as.data.frame(strsplit(names(tapply(all.ev$gfp.rawratio, all.ev$combo_temp, mean)), "_"))[1, ]
mch.levels <- as.data.frame(strsplit(names(tapply(all.ev$gfp.rawratio, all.ev$combo_temp, mean)), "_"))[2, ]
temp.plot <- as.data.frame(strsplit(names(tapply(all.ev$gfp.rawratio, all.ev$combo_temp, mean)), "_"))[3, ]

myPalettePurple <- colorRampPalette(rev(brewer.pal(9, "Purples")) )
myPalettePink <- colorRampPalette((brewer.pal(9, "PiYG")) )

#center community fraction heatmaps at white = 0
#create custom color vector with purples and pinks
palette.vec <- rev(c(myPalettePink(11)[1:5], "white", rev(myPalettePurple(9)[1:7]) ))

## Make 3 panel figure
par(mfrow = c(1, 3))

# Create manual range for 32 and 37 color bar, to make them the same
plot.range <- c(.5, 1.31)
# Create vector of range of values at 42 for color bar
range.ratio42 <- as.numeric(range(as.numeric(name.split42$ratio)))

p1 <- ggplot(name.split32, aes(x=as.numeric(gfp.time), y=as.numeric(mch.time))) + geom_tile(aes(fill = as.numeric(ratio)),colour='white')  + ggtitle("GFP Ratios 32C") + xlab("GFP Timepoint") + ylab("mCherry Timepoint")  + theme_classic() + theme(legend.title = element_blank() ) + scale_x_continuous( breaks = c(1, 5, 9, 13), expand = c(0, 0)) + scale_y_continuous( breaks = c(1, 5, 9, 13), expand = c(0, 0)) + scale_fill_gradientn(colours=rev(palette.vec[5:13]), limits  = plot.range) + theme(axis.text=element_text(size=12), axis.title=element_text(size=14) , plot.title = element_text(size=18))

p2 <- ggplot(name.split37, aes(x=as.numeric(gfp.time), y=as.numeric(mch.time))) + geom_tile(aes(fill = as.numeric(ratio)),colour='white')  + ggtitle("GFP Ratios 37C") + xlab("GFP Timepoint") + ylab("mCherry Timepoint")  + theme_classic() + theme(legend.title = element_blank() ) + scale_x_continuous( breaks = c(1, 5, 9, 13), expand = c(0, 0)) + scale_y_continuous( breaks = c(1, 5, 9, 13), expand = c(0, 0))  + scale_fill_gradientn(colours=rev(palette.vec[5:13]), limits  = plot.range) + theme(axis.text=element_text(size=12), axis.title=element_text(size=14) , plot.title = element_text(size=18))

p3 <- ggplot(name.split42, aes(x=as.numeric(gfp.time), y=as.numeric(mch.time))) + geom_tile(aes(fill = as.numeric(ratio)),colour='white')  + ggtitle("GFP Ratios 42C") + xlab("GFP Timepoint") + ylab("mCherry Timepoint")  + theme_classic() + theme(legend.title = element_blank() ) + scale_x_continuous( breaks = c(1, 5, 9, 13), expand = c(0, 0)) + scale_y_continuous( breaks = c(1, 5, 9, 13), expand = c(0, 0)) + scale_fill_gradientn(colours=rev(palette.vec[c(1,1, 1:8, 11, 13)]), limits  = c(range.ratio42[1]  , range.ratio42[2]  ) ) + theme(axis.text=element_text(size=12), axis.title=element_text(size=14) , plot.title = element_text(size=18))

grid.arrange(p1, p2, p3, nrow = 1)


# Look at whether popualtion size at 42 is related to significant interaction, to evaluate whether abundance could drive an artifact
tapply(all.ev$gfpstd[all.ev$temp == 42], all.ev$lineage[all.ev$temp == 42], mean)

# Lineages with interaction at 42: 1, 2, 7, 10, 11, 13, 15, 18, 19, 20, 21, 22
mean(tapply(all.ev$gfpstd[all.ev$temp == 42], all.ev$lineage[all.ev$temp == 42], mean)[c(1, 2, 7, 10, 11, 13, 15, 18, 19, 20, 21, 22 )])
mean(tapply(all.ev$gfpstd[all.ev$temp == 42], all.ev$lineage[all.ev$temp == 42], mean)[-c(1, 2, 7, 10, 11, 13, 15, 18, 19, 20, 21, 22 )])
sd(tapply(all.ev$gfpstd[all.ev$temp == 42], all.ev$lineage[all.ev$temp == 42], mean)[c(1, 2, 7, 10, 11, 13, 15, 18, 19, 20, 21, 22 )])
sd(tapply(all.ev$gfpstd[all.ev$temp == 42], all.ev$lineage[all.ev$temp == 42], mean)[-c(1, 2, 7, 10, 11, 13, 15, 18, 19, 20, 21, 22 )])

# No significant difference in mean or variance of fraction of population comprised
```

```
## Create script to analyze merged growth rate data

library(lme4)
library(KernSmooth)
library(viridis)
library(RColorBrewer)
library(car)

# Make a function to difference the log OD values
log.diff <- function(vector){
  diffs <- ((vector[-1])) - (vector[-length(vector)])
}

smooth.fun <- function(vec, width) {
  #Find how many values are NA
  nas <- sum(is.na(vec))
  new.length <- length(vec) - nas

  x.vals <- seq(1, new.length)
  y.vals <- log(vec[1:new.length] + .02)

  smoothed.y <- locpoly(x.vals, y.vals, degree = 1, bandwidth = width, gridsize = new.length)$y

  return(smoothed.y)
}

## Read in merged dataset
g.all <- read.csv("~/Desktop/AllGrowthCurves.csv", header = T, row.names = 1)

## Split up merged dataset by temperature and resistance timepoint
c37.all <- g.all[, grepl("C37", colnames(g.all))]
c32.all <- g.all[, grepl("C32", colnames(g.all))]
c42.all <- g.all[, grepl("C42", colnames(g.all))]

# Make smoothed datasets
#### Choose smoothing width
width.all <- 24 # results don't change qualitatively within the range of 12 to 30


c37.mins <- apply(c37.all, 2, min, na.rm = T)
c37.zeroed <- sweep(c37.all, MARGIN = 2, STATS  = c37.mins, FUN = "-")
c37.smooth <- apply(c37.zeroed, 2, smooth.fun, width = width.all)
c37.diffs <- lapply(c37.smooth, log.diff)
all.c37.maxdiffs <- unlist(lapply(c37.diffs, max))

c32.mins <- apply(c32.all, 2, min, na.rm = T)
c32.zeroed <- sweep(c32.all, MARGIN = 2, STATS  = c32.mins, FUN = "-")
c32.smooth <- apply(c32.zeroed, 2, smooth.fun, width = width.all)
c32.diffs <- lapply(c32.smooth, log.diff)
all.c32.maxdiffs <- unlist(lapply(c32.diffs, max))

c42.mins <- apply(c42.all, 2, min, na.rm = T)
c42.zeroed <- sweep(c42.all, MARGIN = 2, STATS  = c42.mins, FUN = "-")
c42.smooth <- apply(c42.zeroed, 2, smooth.fun, width = width.all)
c42.diffs <- lapply(c42.smooth, log.diff)
all.c42.maxdiffs <- unlist(lapply(c42.diffs, max))


## Do same analysis with relative fitness

# Create vectors for anvestral growth rates (at T1)
anc37 <- all.c37.maxdiffs[1:24]
anc32 <- all.c32.maxdiffs[1:24]
anc42 <- all.c42.maxdiffs[1:24]

rel.c37.maxdiffs <- all.c37.maxdiffs / rep(anc37, times = 14)
rel.c32.maxdiffs <- all.c32.maxdiffs / rep(anc32, times = 14)
rel.c42.maxdiffs <- all.c42.maxdiffs / rep(anc42, times = 14)

##Create vector corresponding to resistance timepoint for the above maximum growth rates
res.level <- rep(1:14, each = 24)

## Make plot of average growth rate with std err bars at 3 temperatures, with color changing by resistance level
scale_colour_manual(values = rainbow(8))
par(mfrow = c(1, 2))
plot(0, 0, xlim = c(28, 44), ylim = c(.016, .038), xlab = "Temperature", ylab = "Maximum Growth Rate" , main = "Absolute Growth Rate")
for(tp in 1:14){
  gr32 <- mean(all.c32.maxdiffs[res.level == tp])
  se32 <- sd(all.c32.maxdiffs[res.level == tp]) / sqrt(24)

  gr37 <- mean(all.c37.maxdiffs[res.level == tp])
  se37 <- sd(all.c37.maxdiffs[res.level == tp]) / sqrt(24)

  gr42 <- mean(all.c42.maxdiffs[res.level == tp])
  se42 <- sd(all.c42.maxdiffs[res.level == tp]) / sqrt(24)

  points(c(gr32, gr37, gr42) ~ c(32, 37, 42), type = "l", col = rev(viridis(14))[tp], lwd = 2)
  points(c(gr32, gr37, gr42) ~ c(32, 37, 42), type = "p", col = rev(viridis(14))[tp], pch = 20)
  segments(y0 = c(gr32 + se32/2, gr37 + se37/2, gr42 + se42/2),
           y1 = c(gr32 - se32/2, gr37 - se37/2, gr42 - se42/2),
           x0 = c(32, 37, 42),
           lwd = 3,
           col = rev(viridis(14))[tp])

}

legend("topleft", legend = seq(1:14), pch = 20, col = rev(viridis(14)), x.intersp = .5, bty = "n")

plot(0, 0, xlim = c(30, 44), ylim = c(.59, 1.05), xlab = "Temperature", ylab = "Maximum Growth Rate", main = "Relative Growth Rate" )
for(tp in 1:14){
  gr.rel32 <- mean(rel.c32.maxdiffs[res.level == tp])
  se.rel32 <- sd(rel.c32.maxdiffs[res.level == tp]) / sqrt(24)

  gr.rel37 <- mean(rel.c37.maxdiffs[res.level == tp])
  se.rel37 <- sd(rel.c37.maxdiffs[res.level == tp]) / sqrt(24)

  gr.rel42 <- mean(rel.c42.maxdiffs[res.level == tp])
  se.rel42 <- sd(rel.c42.maxdiffs[res.level == tp]) / sqrt(24)

  points(c(gr.rel32, gr.rel37, gr.rel42) ~ c(32, 37, 42), type = "l", col = rev(viridis(14))[tp], lwd = 2)
  points(c(gr.rel32, gr.rel37, gr.rel42) ~ c(32, 37, 42), type = "p", col = rev(viridis(14))[tp], pch = 20)
  segments(y0 = c(gr.rel32 + se.rel32/2, gr.rel37 + se.rel37/2, gr.rel42 + se.rel42/2),
           y1 = c(gr.rel32 - se.rel32/2, gr.rel37 - se.rel37/2, gr.rel42 - se.rel42/2),
           x0 = c(32, 37, 42),
           lwd = 3,
           col = rev(viridis(14))[tp])

}

## Format data for regression analysis

## Concatenate all growth rate values into one vector
p1.abs <- c( all.c32.maxdiffs, all.c37.maxdiffs, all.c42.maxdiffs)

## Relative fitess: divide growth rates by respective ancestral growth rates
p1 <-  c( rel.c32.maxdiffs, rel.c37.maxdiffs, rel.c42.maxdiffs)

# Create vector for corresponding temps
t1 <- c(rep(c("c32", "c37", "c42"), each = length(p1.abs) / 3) )

# Create vector for corresponding resistance levels
r1 <- rep(rep(1:14, each = 24 ), times = 3 )

# Create vector for lineages
s1 <- rep(1:24, times = length(p1.abs) / 24 )

# Take out any points where strains didn't grow (defined as relative growth rate less than 1% of ancestor)
# The data points with low growth correspond to wells with no change in optical density
p.1 <- p1[p1 > .1]
t.1 <- t1[p1 > .1]
r.1 <- r1[p1 > .1]
s.1 <- as.factor(s1[p1 > .1])

## Count how many strains had increased relative growth in any evolved strain at 37 versus 32 and 42
sum(t1 == "c37" & p1 > 1)
sum(t1 == "c32" & p1 > 1)
sum(t1 == "c42" & p1 > 1)

##############################################################################################################################
# Regression analysis of relative growth rate costs

summary(lmer(p.1 ~ t.1*r.1 + (1|s.1)))
anova(lmer(p.1 ~ t.1*r.1 + (1|s.1)))
summary(lm(p.1 ~ t.1*r.1 + s.1))
anova(lm(p.1 ~ t.1*r.1 + s.1))
```

```
## Script to analyze mutation data and merge with competition event data

library(lme4)

mu <- read.csv("~/Desktop/AnnotatedMutationsWithCategories.csv", header = T)
ev <- read.csv("~/Desktop/MergedFlowCytometryDataset.csv", header = T)

# Create a new matrix for this information, which will then be merged into events dataset
lin.vec <- rep(seq(1, 24, 1), each = 4)
time.vec <- rep(c(1, 5, 9, 13), times = 24)
str.vec <- paste(lin.vec, time.vec, sep = "_")

# Create matrix for aggregate mutation numbers
ag <- cbind(lin.vec, time.vec)
ag <- as.data.frame(ag)
rownames(ag) <- str.vec
colnames(ag) <- c("Lineage", "Time")

# Create matching name in mu matric
mu$match.name <- paste(mu$Lineage, mu$Time, sep = "_")

# Create matching name format in events dataframe
ev$gfp.name <- sub('.', '', ev$gfpstrain)
ev$mch.name <- sub('.', '', ev$mchstrain)

# Create a function to find the immediate ancestor of a strain
find.ancestor <- function(strain.name){
  lineage <- as.numeric(unlist(strsplit(strain.name, "_"))[1])
  timepoint <- as.numeric(unlist(strsplit(strain.name, "_"))[2])

  ifelse(timepoint == 1,
         name.return <- NA,
         name.return <- paste(lineage, time.vec[which(unique(time.vec) == timepoint) - 1], sep = "_")
  )

  return(name.return)
}

find.ancestor("1_9")


## For counting categories of mutations, create logical column for whether strain has the mutation, and then count them
# Create column for each category of gene

for(cat in unique(mu$GeneName)){
  cat.vec <- mu$GeneName == cat
  num.cat <- tapply(cat.vec, mu$match.name, sum)

  ag[, get("cat")] <- num.cat[match(rownames(ag), names(num.cat))]
  ag[, get("cat")][ ! rownames(ag) %in% names(num.cat)] <- 0

}

## Make columns in events dataframe for difference in mutations between competed strains
# Find matching values for each strain, subtract them, and create new column named "DiffIn__"

col.compare <- colnames(ag)[-c(1, 2)]

for(col.create in col.compare){
  str1.val <- ag[, get("col.create")][match(ev$gfp.name , rownames(ag))]
  str2.val <- ag[, get("col.create")][match(ev$mch.name , rownames(ag))]
  new.col.name <- paste0("DiffIn", get("col.create"))


  ev[, get("new.col.name")] <- str1.val - str2.val
}


# Relevel to use 37 as the standard so the linear model compares against this temp
ev$temp <- as.factor(ev$temp)
ev$temp <- relevel(ev$temp, ref = "37")

# Remove duplicate columns for genes with same mutation profile
duplicated(as.list(ev))
colnames(ev)[duplicated(as.list(ev))]
ev <- ev[!duplicated(as.list(ev))]


lm.genes <- summary(lm(log(ev$gfp.rawratio) ~
                           ev$DiffInyahF * ev$temp +
                           ev$DiffInaraD * ev$temp +
                           ev$DiffInyaiO * ev$temp +
                           ev$DiffInacrB * ev$temp +
                           ev$DiffInacrA * ev$temp +
                           ev$DiffInacrR * ev$temp +
                           ev$DiffInacrR.mscK * ev$temp +
                           ev$DiffInmscK * ev$temp +
                           ev$DiffInmdfA* ev$temp +
                           ev$DiffInompF * ev$temp +
                           ev$DiffInopgH * ev$temp +
                           ev$DiffInicd.ymfD * ev$temp +
                           ev$DiffInlomR * ev$temp +
                           ev$DiffInpaaC * ev$temp +
                           ev$DiffIndgcF * ev$temp +
                           ev$DiffInmarR * ev$temp +
                           ev$DiffInmarA * ev$temp +
                           ev$DiffInmarB * ev$temp +
                           ev$DiffInyejA * ev$temp +
                           ev$DiffInyfaQ.yfaT * ev$temp +
                           ev$DiffInmenH * ev$temp +
                           ev$DiffInxdhB * ev$temp +
                           ev$DiffInagaS * ev$temp +
                           ev$DiffInelbB.arcZ * ev$temp +
                           ev$DiffInarcB * ev$temp +
                           ev$DiffInrplM * ev$temp +
                           ev$DiffInenvZ * ev$temp +
                           ev$DiffInompR * ev$temp +
                           ev$DiffInatpC * ev$temp +
                           ev$DiffInatpD * ev$temp +
                           ev$DiffInatpA * ev$temp +
                           ev$DiffInatpH * ev$temp +
                           ev$DiffInatpF * ev$temp +
                           ev$DiffInatpE * ev$temp +
                           ev$DiffInatpB * ev$temp +
                           ev$DiffInuvrA * ev$temp +
                           ev$DiffInuvrA.ssb * ev$temp +
                           ev$DiffInssb * ev$temp +
                           ev$DiffInuxuR * ev$temp +
                           ev$DiffInslt * ev$temp +
                           ev$DiffIncreB * ev$temp +
                           ev$lineage * ev$temp ) )

# Convert coefficients into dataframe
gene.coef <- lm.genes$coefficients
lm.genes

## Count significant differences in genes

# Find effect sizes for genes at different temperatures
par(mfrow = c(2, 1))
eff32 <- gene.coef[, 1][grepl("Diff", rownames(gene.coef)) & grepl("temp32", rownames(gene.coef)) ]
eff37 <- gene.coef[, 1][grepl("Diff", rownames(gene.coef)) & !grepl("temp32", rownames(gene.coef)) & !grepl("temp42", rownames(gene.coef)) ]
eff42 <- gene.coef[, 1][grepl("Diff", rownames(gene.coef)) & grepl("temp42", rownames(gene.coef)) ]

par(mfrow = c(3, 1))
hist(eff37 + eff32, breaks = 10, xlim = c(-2.5, 2.5), ylim = c(0, 10), main = "Estimated Gene Effects at 32 Degrees", xlab = "Effect Size")
#abline(v = median(eff37 + eff32))
hist(eff37 , breaks = 10, xlim = c(-2.5, 2.5), ylim = c(0, 10), main = "Estimated Gene Effects at 37 Degrees", xlab = "Effect Size")
#abline(v = median(eff37 ))
hist(eff37 + eff42, breaks = 15, xlim = c(-2.5, 2.5),ylim = c(0, 10),  main = "Estimated Gene Effects at 42 Degrees", xlab = "Effect Size")
#abline(v = median(eff37 + eff42))

effect.32 <- eff37 + eff32
effect.42 <- eff37 + eff42
var.test(effect.32, eff37, alternative = "two.sided")
var.test(effect.42, eff37, alternative = "two.sided")

####
#### Make barchart for mutations at different time points and analyze distribution of categories over time

## Make a plot of new mutations arising at different time points

new.mu <- mu[mu$Time == 1, ]

for(str in unique(mu$match.name)){
  if(unlist(strsplit(str, "_"))[2] > 1){
    anc <- find.ancestor(str)

    str.mut <- mu[mu$match.name == str, ]
    anc.mut <- mu[mu$match.name == anc, ]

    new.mu <- rbind(new.mu, str.mut[!str.mut$Location %in% anc.mut$Location, ])

  }
}


# Count mutations in each category at each time point
tapply(new.mu$Category2[new.mu$Time == 1], new.mu$Category2[new.mu$Time == 1], length)
tapply(new.mu$Category2[new.mu$Time == 5], new.mu$Category2[new.mu$Time == 5], length)
tapply(new.mu$Category2[new.mu$Time == 9], new.mu$Category2[new.mu$Time == 9], length)
tapply(new.mu$Category2[new.mu$Time == 13], new.mu$Category2[new.mu$Time == 13], length)

# Create dataset for barchart
new.counts <- c(
  tapply(new.mu$Category2[new.mu$Time == 1], new.mu$Category2[new.mu$Time == 1], length),
  tapply(new.mu$Category2[new.mu$Time == 5], new.mu$Category2[new.mu$Time == 5], length),
  tapply(new.mu$Category2[new.mu$Time == 9], new.mu$Category2[new.mu$Time == 9], length),
  tapply(new.mu$Category2[new.mu$Time == 13], new.mu$Category2[new.mu$Time == 13], length)
)

#Create times for barchart
chart.times <- as.factor(c(
  rep(1, times = length(tapply(new.mu$Category2[new.mu$Time == 1], new.mu$Category2[new.mu$Time == 1], length) ) ),
  rep(5, times = length(tapply(new.mu$Category2[new.mu$Time == 5], new.mu$Category2[new.mu$Time == 5], length) ) ),
  rep(9, times = length(tapply(new.mu$Category2[new.mu$Time == 9], new.mu$Category2[new.mu$Time == 9], length) ) ),
  rep(13, times = length(tapply(new.mu$Category2[new.mu$Time == 13], new.mu$Category2[new.mu$Time == 13], length) ) )
) )

Category <- names(new.counts)

bar.chart <- data.frame(new.counts, Category, chart.times)

## Re-order bar chart to have colors go by mean appearance time
bar.chart$time.mult <- bar.chart$new.counts * as.numeric(bar.chart$chart.times)
mean.times <- tapply(bar.chart$time.mult, bar.chart$Category, sum) / tapply(bar.chart$new.counts, bar.chart$Category, sum)
bar.chart$Category <- factor(bar.chart$Category, levels = rev(names(sort(mean.times)) ) )

ggplot(bar.chart, aes(fill=Category, y=new.counts, x=chart.times)) +
  geom_bar(position="stack", stat="identity") +
  xlab("Timepoint") +
  ylab("Number of New Mutations")

## Now, chi-squared tests of distribution of mutations over time
# Evaluate which categories are unevenly distributed among timepoints

chi.sq <- vector()
for(cat2 in unique(bar.chart$Category)){
  place <- which(cat2 == unique(bar.chart$Category))
  chi.sq[place] <- (chisq.test(bar.chart$new.counts[bar.chart$Category == cat2]))$p.value
}

names(chi.sq) <- unique(bar.chart$Category)

# Count number of mutations at different times and in different categories
tapply(mu$Time, mu$Time, length)
tapply(mu$Category2, mu$Category2, length)
tapply(new.mu$Time, new.mu$Time, length)

## Check whether any specific mutation category is associated with the presence of greater cost at 42 degrees
# Lineages with interaction at 42: 1, 2, 7, 10, 11, 13, 15, 18, 19, 20, 21, 22

# Create matrix for presence or absence of each mutation category in each lineage
lin.cat <- vector()
for(c1 in unique(mu$Category2)){
  lin.tf <- as.numeric(seq(1:24) %in% unique(mu$Lineage[mu$Category2 == c1]))
  lin.cat <- rbind(lin.cat, lin.tf)
}

rownames(lin.cat) <- unique(mu$Category2)
colnames(lin.cat) <- 1:24

# Do Fisher's exact test on lineages with interactions and mutation category
int.lins <- c(1, 1, 0, 0, 0, 0, 1, 0, 0, 1, 1, 0, 1, 0, 1, 0, 0, 1, 1, 1, 1, 1, 0, 0)
names(int.lins) <- seq(1:24)

apply(lin.cat, 1, fisher.test, y = int.lins, simulate.p.value = T)

# Calculate probability of finding 5 significant categories by chance
# Equals 10 choose 5 (252) * .05^5 * .95^5
252 * (.05^5) * .95^5
```
